# Micro-to-Macro Scale Hydrogel Microchannel Networks by Twisted Wire Templating

**DOI:** 10.64898/2026.03.24.713957

**Authors:** Jiayu Deng, Weizhen Pan, Fahmida Alom, Hassan Tahir, Yifeng Xuan, Bian Lijing, Brian Cunningham, Sam H. Au

**Affiliations:** Department of Bioengineering, Imperial College London; London, SW7 2AZ, United Kingdom; Cancer Research UK Convergence Science Centre; London, SW7 2AZ, United Kingdom

**Keywords:** microfluidics, microvasculature, multiscale models, hierarchical bifurcations, wire twisting

## Abstract

The human vasculature is a complex, multiscale system comprising hierarchical networks of macroscale to microscopic vessels. Existing in vitro fabrication techniques often fail to bridge these disparate scales, as high-resolution methods like multiphoton ablation are too slow for replicating larger vessels, while 3D printing lacks the resolution for fine microscale features. Here, we report a “twisted wire templating” strategy capable of generating perfusable bifurcating hydrogel networks that seamlessly transition from the macro- to the micro-scale (2.3 mm to 140 µm) through seven orders of bifurcations. By optimizing wire-twisting geometries and polyurethane dip-coating, we overcame instability-driven bead formation to ensure replication fidelity across the networks. Fabrication rigs were reconfigured from existing 2D planar layouts to 3D reconfigurable architectures to better replicate 3D vessel geometries which simultaneously reducing the laboratory footprint and fabrication times by 47%. Using a Taguchi orthogonal array, we further optimized surface chemistry and hydrogel composition to inhibit structural failure during template extraction, resulting in fully patent, perfusable networks. This method provides a robust, low-cost, and scalable foundation for creating physiologically representative vascular models for investigating multiscale disease mechanisms and organ-level tissue engineering.

## 1 Introduction

Blood vessels are organised into fractally bifurcating networks that span several length-scales from macroscale arteries to microscopic microvasculature and capillaries. This multiscale hierarchy spanning these scales each present with unique structure, function, and hemodynamics^1^. These unique characteristics across blood vessels scales have important implications in the pathophysiology of many diseases. For instance, changes in blood vessel diameters force circulating tumour cell clusters to dynamically reorganise their shape as they navigate through the circulation^2^, inflammatory signals released by microscale blood vessels promotes coronary artery diseases and atherosclerotic plaque formation in larger vessels^3^ and the narrowing of large cerebral vessels contributes to the development of Alzheimer’s and other neurodegenerative diseases by inducing dysfunction in capillaries^4^.

High fidelity *in vitro* models of human vasculature that spans micro to macro scales are needed to investigate disease mechanisms and to refine and advance therapies. Microfluidics has emerged as a promising technology for mimicking the microenvironments of human blood vessels due to their ability to precisely control fluid flows, vascular geometry and multicellular interactions without the need for animal experimentation^5,6^. Some techniques, such as 3D printing are well suited to generating macroscale vessel models through direct or sacrificial moulding, but have resolution limits that constrain the production of narrow microscale networks^7,8^. Other fabrication methods, such as multi-photon ablation are capable of generating microchannels as small as 2-7.5µm^9,10^. However, because this technique operates at the voxel level, it is prohibitively time consuming for developing large-scale vasculature^11^. To date, no single fabrication method has successfully generated perfusable networks of bifurcating microchannels from micro (i.e. micrometer) to macro (i.e. multi-millimetre).

Our group has recently developed a surface-tension driven wire templating method able to generate perfusable bifurcating microchannel networks that fractally narrow down to capillary scale (6.1±0.3 µm)^12^. We have thus far demonstrated the ability of this technique to produce two orders of bifurcations (i.e. 4 circular cross-section channels into 2 into 1). Because of low cost and the repeatable nature of generating bifurcating wires using surface-tension, this method may be uniquely capable of generating higher orders of micro-to-macroscale vessels. However, to achieve this goal, two main complications must be addressed (Figure 1): (a) the method of generating wire template networks is labour intensive and slow, taking an estimated 6.5 hours to generate a 7 order (i.e. 128 - 64 - 32 - 16 - 8 - 4 - 2 - 1) network wire template, (b) the planar 2D geometries of existing dipping and alignment rigs necessary for generating wire templates and casting them into hydrogels are poorly representative of human vessel network sand would be unwieldy if simply scaled up.

**Figure 1.**
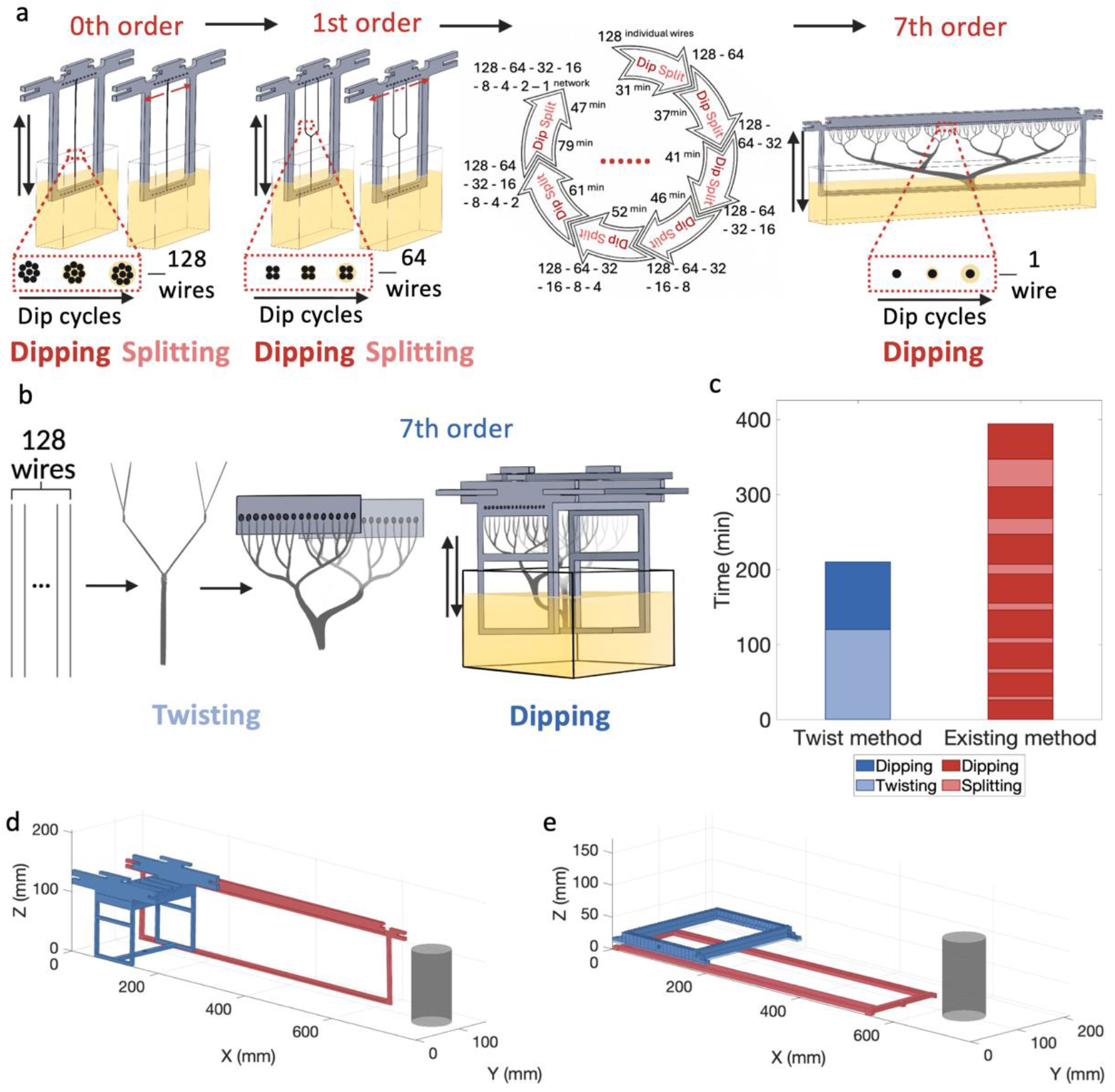
Comparison of the existing and twisting wire template fabrication workflows and rig architectures. (a) Existing method including sequential dipping and splitting steps needed to generate 7 order networks. (b) Twist method fabrication workflow involves such one dipping cycle. (c) Comparison of fabrication time between the two methods. Dimensional comparison of the existing (red) and updated (blue) (d) dipper rigs and (e) alignment rigs. A standard 350 mL can (grey) is shown for scale.

In this work, we describe modifications to our technique that involve twisting of individual template wires to simplify the generation of higher order wire templates by condensing sequential dipping processes into a single dip-coating step that reduces total fabrication time by 47% alongside transitioning from 2D planar to 3D layouts that represent more physiological arrangements whilst reducing the footprints of the dipping and alignment rigs to make them compatible with typical laboratory benches. This approach allowed for the rapid generation of perfusable networks consisting of seven orders of bifurcations and up to 128 individual microchannels with dimensions of ∼100 microns.

## 2 Results

### 2.1 Addressing Scalability in High-Order Vascular Networks

The existing method of fabricating wire template networks involves a successive dip-coating process where wires are: strung up on an alignment rig, dip coated in a bath of polyurethane (PU), separated and re-strung to initiate a bifurcation, and then re-dripped for each branch (Figure 1a). While effective for low-order networks, we found that scaling this process to a seventh order network consisting of 128 individual wires was poorly scalable. We estimate that a skilled operator would require approximately 6.5 hours without break to generate a seventh order wire template. To remove the need for successive coating and manual wire splitting steps, we developed a strategy of twisting wire bundles prior to dipping which reduced the necessary dipping cycles to one.

Another challenge we encountered when fabricating higher order networks is that the physical size of existing rigs used to assist dipping and hydrogel casting became unwieldly. This was because previous rigs were designed to array wires in 2D planar geometries; scaling to a seven-order bifurcation would result in a dipper rig width of 645mm and alignment rig width of 553mm (Figure 1d and 1e). This is impractical for use in standard laboratory settings and fails to reflect the layout of a 3D dimensional network of high-order networks. To overcome these spatial constraints, we aimed to transition from 2D planar layouts to 3D configurations. By ‘bundling’ the branching wires in three dimensions, we hoped to condense the dipper and alignment rig widths to 217mm and 210mm, respectively.

### 2.2 Development of wire twisting protocol for bifurcations

To reduce the labour and time required for fabrication of high-order wire template networks, we explored a strategy of twisting wires near bifurcations to reduce the labour of wire template generation, while reducing the number of PU dips required to one. Three twisting configurations were evaluated: twist-at-below, all twist, and twist-at-both (Figure 2a–c).

**Figure 2.**
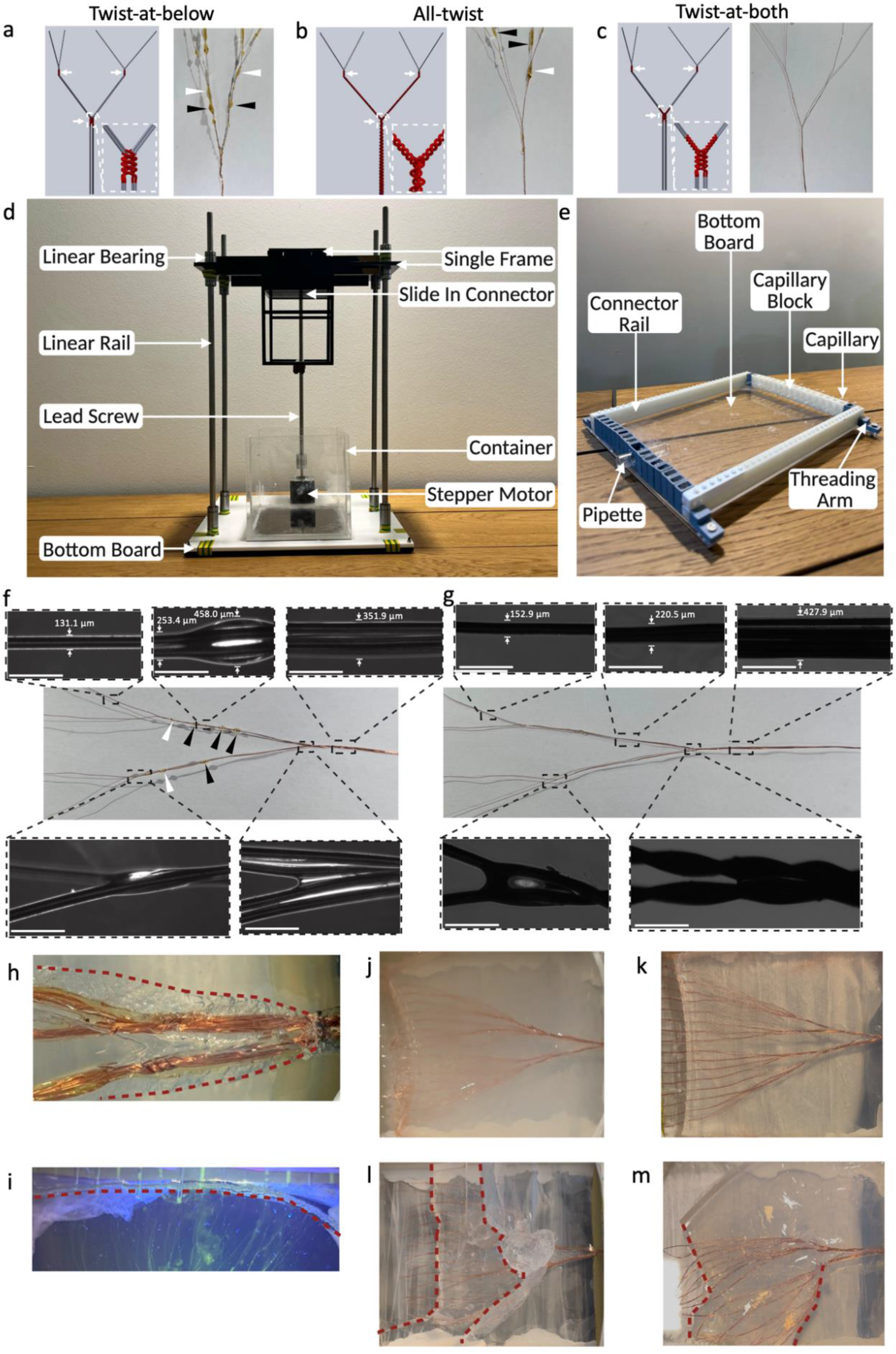
Optimisation of twisting strategy, rig redesign, dip-coating parameters, and orthogonal test conditions for seven-order wire template and hydrogel channel fabrication. Three twisting strategies for bifurcation nodes: (a) twist-at-below, (b) all-twist, and (c) twist-at-both, shown as schematics (left) and images of representative PU-coated wire templates (right). White arrows indicate twisted regions, white arrowheads indicate beads at bifurcation, and black arrowheads indicates beads at branches. (d) Re-designed 3D dipper rig. (e) Re-designed modular alignment rig. Dip-coating cycle optimisation at 1 mm s^−1^ using two-order bifurcating wire templates: (f) 25 dipping cycles, (g) 15 dipping cycles. Scale bar: 50 μm. White arrowheads indicate beads at bifurcation, and black arrowheads indicates beads at branches. Polyacrylamide hydrogels generated from wire templates showing representative images of crack formation near (h) the inlet hydrogel with wire template inside and (i) outlet of FITC-dextran perfused channel networks after wire extraction. Red dashed lines indicate cracked regions. Representative images of hydrogel with embedded wire template from orthogonal test results with various fabrication settings: (j) no TMSPMA capillary coating, PU wire template coating, 5% (w/v) acrylamide; (k) no TMSPMA capillary coating, PU and TMSCl wire template coating, 7% (w/v) acrylamide; (l) with TMSPMA capillary coating, PU wire template coating, 7% (w/v) acrylamide; (m) with TMSPMA capillary coating, PU and TMSCl wire template coating, 5% (w/v) acrylamide. Red dashed lines indicate cracked regions.

In the twist-at-below configuration, the wires were twisted only after the daughter branches met at the bifurcation node (Figure 2a). Although this configuration preserved the overall bifurcation geometry, pronounced PU bead formation was observed during dip-coating. Beads were frequently located near the node region (Fig. 2a, white arrowhead) and along the parental branches (Fig. 2a, black arrowhead), resulting in an uneven coating profile.

In contrast, the all-twist configuration involved twisting the entire wire length both upstream and downstream of the bifurcation (Figure 2b). Although this configuration improved the mechanical stability of the wire bundles, pronounced bead formation was observed near the bifurcation nodes (Figure 2b, white arrowheads), and the coating thickness varied markedly along the twisted segments (Figure 2b, black arrowheads).

The third strategy, twist-at-both, introduced short, twisted regions on both sides of each bifurcation node (Figure 2c). This configuration produced wire templates in which the branches remained tightly coupled at the bifurcation and the coating was most uniform between nodes. Compared with the other two strategies, substantially reduced PU beads formation, and the coated surface appeared smoother across both twisted and non-twisted regions.

Based on these observations, the twist-at-both strategy was selected for subsequent experiments fabricating higher-order bifurcating wire templates.

### 2.3 Re-engineering of the Rig System for High-Order Bifurcation

To overcome the limited bifurcation order and spatial constraints of the previous planar system, both the dipper and alignment rigs were re-engineered into 3D configurations. In the redesigned dipper rig (Figure 2d), the first three bifurcations align in the y-z plane, with subsequent branches distributed in the x-y plane, featuring 128 slots to simultaneously load a full seven-order wire template network. Structurally, a single upper frame provides mechanical stability for the detachable slide-in connectors, which modularise and streamline the loading of wire bundles. To execute precise vertical dipping motions, the rig is equipped with a stepper motor and a lead screw, guided smoothly by linear rails and bearings along the z-axis into the bottom container.

Similarly, the redesigned alignment rig utilises a modular interlocking approach to support more complex wire templates (Figure 2e). The reconfigurable assembly consists of capillary blocks to hold individual capillaries in precise alignment, threading arms to anchor securing threads, and connector rails that link these elements into a continuous frame upon a stabilising bottom board. Each capillary block was designed to house a single guide glass capillary into which a bundle of eight of the smallest dimension wires are inserted. After loading, the bundle was pushed back to allow the individual wires to separate, thereby generating the final bifurcation order and forming individually addressable terminal channels. This allows for eight-fold fewer guide glass capillaries to be used within a more compact footprint. Additionally, a pipette is integrated at the terminal end to hold the thickest end of the wire template before the bifurcation starts. Together, these components form a highly adaptable system that can be rapidly adjusted to meet the specific bifurcation requirements of diverse wire templates.

### 2.4 Optimisation of dip-coating parameters

Our previous work identified that dip coating parameters such as dipping velocity and number of dips were important parameters for controlling the uniformity, thickness, and regularity of PU coated wire templates. Prior to evaluating the effects of dipping cycles, we therefore optimised the dipping velocity (the linear velocity at which templates wires were dipped into and removed from PU baths) using two-order wire templates generated on dipping rigs (Fig. 2d). Detailed velocity optimisation results are provided in the Supplementary Figure S1. From these results, a velocity of 1.0 mm s^−1^ was selected as it provided stable coating with minimal bead formation while maintaining fabrication efficiency.

We started with 25-cycles of dips at a 1.0 mm s-1 dip velocity based on previous experience with dip coating. The coated wire template exhibited the expected hierarchical reduction in diameter throughout the bifurcating structure The straight segment before the first bifurcation measured 352μm in diameter, decreasing to 253μm between the first and second bifurcations, and further reduced to 131μm after the second-order bifurcation (Figure 2f). However, extensive bead formation was observed along the branches. Most beads occurred between the first and second bifurcations (Fig. 2f black arrowheads), with bead diameters up to 458μm, nearly 1.8 times larger than the local wire diameter. These occurred despite using the twist-at-both twisting strategies. Beads were also observed near second bifurcation nodes (Fig. 2f white arrowheads), suggesting that excessive dipping cycles promoted PU accumulation at bifurcation regions. These beads introduced surface irregularities, disrupted bifurcation geometry, and may increase the risk of crack formation during subsequent wire extraction.

We therefore conducted additional tests with 15-cycles of dipping instead. These coated wire templates exhibited a smooth and continuous coating along the bifurcating structure (Figure 2g). The diameter decreased from 428µm to 221µm after the first bifurcation and 153µm after the second, demonstrating consistent hierarchical scaling. In contrast to the 25-cycle condition, no significant bead formation was observed along either straight or twisted regions. Detailed dipping cycle optimisation results are provided in the Supplementary Figure S2.

Therefore, 15 dipping cycles at 1 mm s^−1^ were selected as for subsequent fabrication of higher-order bifurcating wire templates.

### 2.5 Optimisation of seven order microchannel network fabrication

We then loaded seven-order PU-coated wire template networks onto alignment rigs (Fig. 2e). Polyacrylamide was poured into the setup and allowed to cross-link, followed by the extraction of wire templates to generate hollow networks. During extraction, we noticed the formation of cracks at opposing ends of the alignment rig: one at the first-order bifurcation (Figure 2h, red dashed line) and another at the glass capillary outlets (Figure 2i, red dashed line). We suspected that the first crack was caused by two factors: (1) friction between the PU-coated wire templates and hydrogel channel wall during extraction, and (2) the high rigidity of the Y-shaped wire template structure at the first-order bifurcation which limited its ability to deform. To address this first type of cracking near the first order bifurcation, we explored surface modification of PU-treated wire template networks with chloro(trimethyl)silane (TMSCl) to increase PU surface hydrophobicity and thereby reduce friction during extraction^18^. In addition, we investigated whether adjusting the concentration of acrylamide could reduce crack formation by altering the mechanical properties like stiffness and stretchability of the hydrogel^19,20^.

For the second crack, we suspected it arose from tensile stress generated by two opposing forces during wire extraction: the strong covalent adhesion between the 3-(trimethoxysilyl) propyl methacrylate (TMSPMA)-coated guide glass capillary and the polyacrylamide hydrogel on one side^21^, and the opposing friction force between the PU surface and the hydrogel channel wall. We therefore explored whether omitting the TMSPMA coating on the guide glass capillary outlets could potentially reduce the formation of the second type of cracks whilst still enabling perfusion at low pressures.

To evaluate the effects of these three factors: capillary coating, wire template surface coating, and acrylamide concentration, a Taguchi L4 orthogonal array (OA) design was employed^22^.

This approach allowed the assessment of the three factors at two levels each (Supplementary Table S1), reducing the number of experimental trials from 8 (2^3^) to 4. Data on the number of first type of cracks and the rate of second-crack formation were collected as response variables to quantify the influence of each factor on crack development (Supplementary Table S3). Following wire extraction, some hydrogels recovered their original structure, whereas others experienced permanent structural failure due to severe cracking. Consequently, a binary outcome, indicating whether the hydrogel network recovered or broke, was also recorded. Representative images of the hydrogels at the point of maximum deformation (Figure 2j, 2k) or failure (Figure 2l, 2m) are presented.

Results were analysed using range analysis, analysis of variance (ANOVA), and regression modelling to compare mean responses between levels and estimate the main effects for each response variable. The combination of no TMSPMS capillary coating, PU with TMSCl template coating, and 5% (w/v) acrylamide was found to minimise crack formation.

### 2.6 Fabrication of seven-order bifurcating wire template and microchannel

Using the optimised coating parameters (dipping velocity of 1mm s^−1^, 15 cycles), we successfully fabricated seven-order, 3D bifurcating wire templates from 128 individual copper wires (100 µm diameter). Macroscopically, the structure demonstrated smooth, continuous PU deposition without bead formation (Figure 3b). Although the 3D architecture naturally obscures some out-of-plane branches in a 2D photograph, the visible bifurcations (Figure 3b, white arrowheads) remained well-aligned. Notably, both the pre- and post-bifurcation twisted regions (Figure 3a) exhibited uniform coating, indicating that the twisting process used to stabilise the nodes did not introduce irregular deformations.

**Figure 3.**
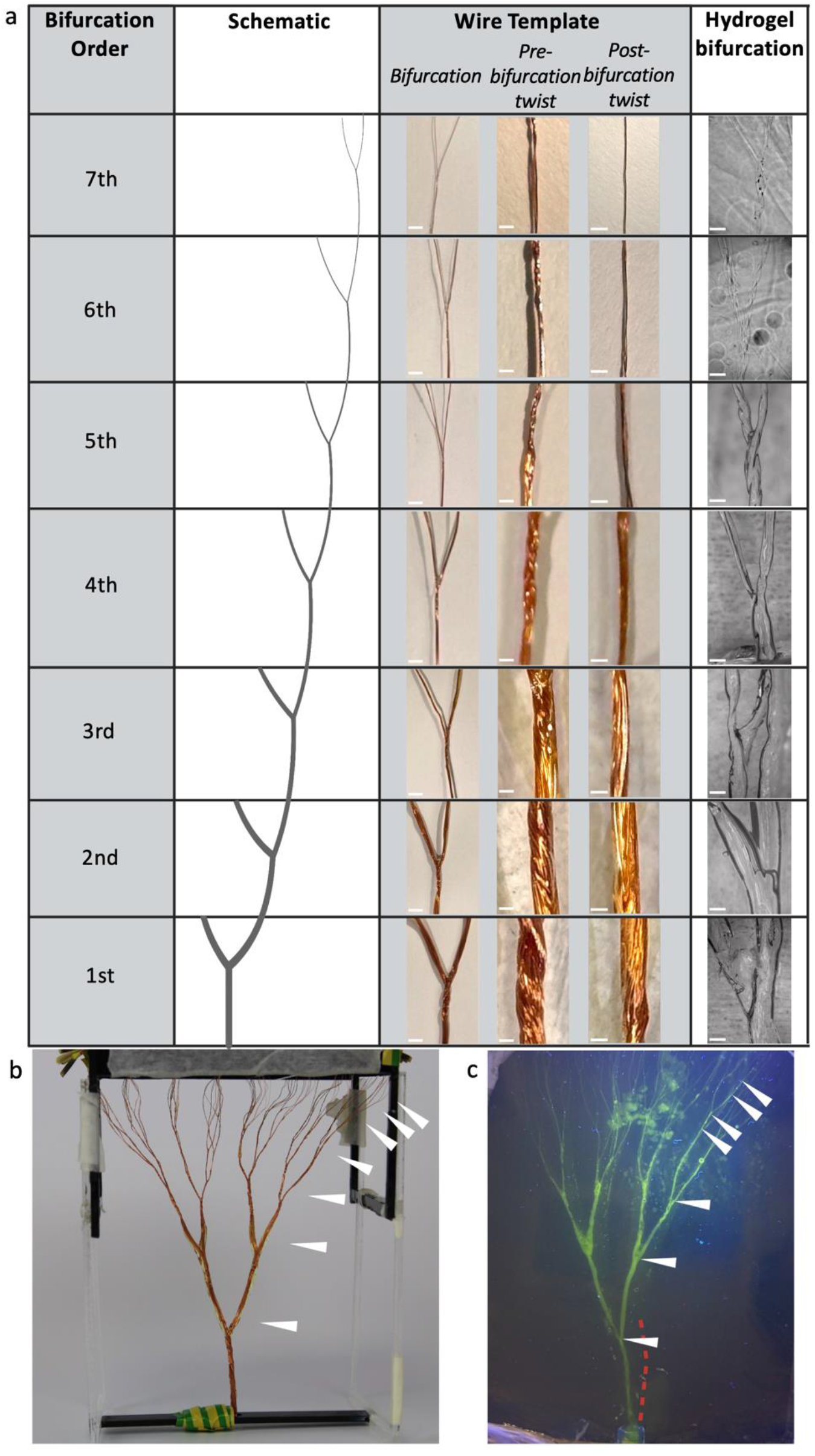
Seven-order bifurcating wire templates and hydrogel channels. (a) Representative images of the wire template and corresponding hydrogel bifurcations from the 1st to 7th order, shown with the schematic, bifurcation region, pre-bifurcation twist, post-bifurcation twist at wire template, and hydrogel bifurcation. Scale bar: 5 mm in the wire template bifurcation column; 1 mm in the pre-bifurcation twist wire template, post-bifurcation twist at wire template, and hydrogel bifurcation columns. (b) Macroscopic view of the seven-order wire template. White arrowheads indicate bifurcation nodes. (c) Macroscopic view of the corresponding seven-order hydrogel channel network after FITC-dextran perfusion. White arrowheads indicate bifurcation nodes, and the red dashed line indicates a representative crack.

Microscopic observations further validated this structural uniformity (Figure 3a). The coated wire diameters transitioned continuously from 1987 ± 114 μm at the primary trunk down to 137 ± 3 μm at the terminal branches (Supplementary Figure S3). These results confirm that our twisting and dip-coating strategy reliably produces geometrically consistent, high-order hierarchical templates.

These optimised templates were subsequently used to cast seven-order polyacrylamide microchannel networks (Figure 3c). Based on our orthogonal testing, we selected a PU + TMSCl template coating and 5% (w/v) acrylamide, while entirely omitting the TMSPMA capillary coating. Notably, even without TMSPMA treatment, fluorescent dye perfusion confirmed that the resulting multi-scale hydrogel network remained fully patent across all 3D branches (Figure 2i, Figure 3c). The overall structural integrity was robust; only one minor crack was observed near the first-order bifurcation region (Figure 3c, red dashed line), and no cracking was detected at the glass capillary outlets.

Finally, the hydrogel channels demonstrated high dimensional fidelity to the wire templates (Figure 3a, right column). Channel diameters seamlessly decreased from 2270 ± 174μm at the inlet to 140 ± 26μm near the outlets (Supplementary Figure S3). Across all seven bifurcation orders, the average absolute dimensional deviation between the template and the hydrogel channels was 9.7 ± 8.9%, confirming the successful translation of macro-to-microscale continuous features.

## 3. Discussion

In this study, we developed a wire twisting method for fabricating higher order bifurcating wire templates, re-designed dipper and alignment rigs from 2D planar to 3D formats, optimised key dip-coating parameters, and refined surface modification and hydrogel parameters using a Taguchi orthogonal array design. Ultimately, the continuous seven-order wire template enabled the formation of perfusable seven-order bifurcating hydrogel channel networks spanning dimensions from 2.3mm to 140μm - mimicking the macro-to-micro vascular structure.

Central to the successful fabrication of these continuous templates was the optimisation of the wire twisting geometry. We speculate that the differences in coating quality among the three tested twisting configurations were primarily governed by how each geometry influenced PU drainage during dip-coating. In the twist-at-below configuration, the relatively large gaps between adjacent parallel wires near the parental branch region (Figure 2a, white arrows) may have promoted local PU accumulation during withdrawal, increasing susceptibility to Plateau-Rayleigh instability and resulting in localised thickening and bead formation^25^. Similarly, in the all-twist configuration, the helical surface generated by continuous twisting (Figure 2b, red twisted regions) likely further inhibited the drainage of excess PU, leading to non-uniform coating thickness and bead formation near both the bifurcation nodes and along the twisted segments. By contrast, the twist-at-configuration emerged as the most favourable geometry for stable coating. The short, twisted regions effectively reduced gap spacing at the bifurcations while preserving relatively smooth and straight segments between nodes. This combination facilitated more uniform PU redistribution during dip-coating, thereby minimising susceptibility to instability-driven bead formation and ensuring excellent overall coating uniformity.

Beyond the high coating uniformity achieved by this optimised geometry, another important advantage of the system developed in this study is its accessibility and low cost. The dipper rig and alignment rig were designed to be fabricated using widely available manufacturing techniques, namely laser cutting and 3D printing, allowing the system to be reproduced with minimal reliance on specialised microfabrication facilities. The rigs were composed of inexpensive materials and most structural elements including higher order wire templates could be reused across multiple experiments.

While recent advancements in fabricating multi-order microvascular networks have increasingly relied on extrusion-based 3D bioprinting of sacrificial templates (such as carbohydrate glass, Pluronic F127, or polyvinyl alcohol^13,14^), these approaches present distinct trade-offs. Additive manufacturing offers extensive geometric freedom, yet it frequently struggles to smoothly transition across multiple length scales. The layer-by-layer deposition process inherently introduces surface roughness or stepped artifacts, particularly along the z-axis, which can perturb local fluid dynamics and induce unnatural physiological shear stresses^15^. Furthermore, water-soluble extruded templates are highly susceptible to swelling and geometric distortion upon contact with aqueous hydrogel precursors prior to complete crosslinking^16^. In contrast, the wire-templated approach presented here circumvents these limitations. By utilising a rigid metallic core and PU coating, the templates resist hydrogel-induced deformation, providing high dimensional fidelity across all seven bifurcation orders. Moreover, the PU dip-coating process naturally yields smooth continuous transitions at the bifurcation nodes that bypass surface irregularities associated with 3D printing. This approach enables the robust construction of high-order hierarchical architectures at a fraction of the cost of high-end bioprinters.

Another key feature of the platform is its modular and flexible design. The rig components can be easily assembled, replaced, or reconfigured, enabling rapid setup and straightforward modification of the system. This modularity allows the platform to accommodate wire templates with different bifurcation orders and geometries. By adjusting the arrangement of modular units such as capillary blocks and connector rails within alignment rigs or selecting wires with different diameters, templates with varying structural dimensions can be generated at will. The transition of dipper and alignment rigs into 3D also enables the fabrication of more anatomically representative models that better reflect the complex, omnidirectional distribution of human vasculature. Together, these features provide a simple and adaptable platform for the fabrication of hierarchical wire templates.

The twisting wire templating technique does have some limitations. First, there is a limited control over the dimensions of the bifurcated wire template, with the optimised technique unable to produce hydrogel networks that precisely follow Murray’s law^23^. Building on previous work^12^, every dip cycle progressively shifted the parent-to-daughter branch diameter ratio closer to the theoretical ratio of symmetrical branching predicted by Murray’s law^24^. The increase in the number of dips, however, leads to progressive thickening of wires, and our data suggests that it may also promote bead formation due to Plateau-Rayleigh instability^25^. According to Landau-Levich-Derjaguin (LLD) theory, reducing the capillary number (Ca) yields thinner PU coatings, enabling finer dimensional control of wire templates^26^. At given wire bundle diameter, dipping velocity is the most obvious parameter for controlling Ca. As we observed, lower dipping velocities led to a reduction in beading likely due to reduced thickness of films, but this must be balanced by longer fabrication times. In future work, the influence of viscosity should also be explored given its influence on Ca. Lower viscosity polymers may promotes faster drainage and reduce the probability of bead formation^27,28^. This may potentially allow more dip cycles to reach Murray’s law before bead instability occurs.

Crack formation during extraction in larger order wire templates was another challenge we encountered. Although the orthogonal test design reduced the rate of crack formation by applying TMSCl to PU-coated wire templates and removing TMSPMA to minimise the tensile stress, we still sometimes encountered cracking near the first-order bifurcation. Our results suggest that while reducing the thickness of PU may mitigate the rigidity of wire templates, doing so compromises the ability of the template to approximate Murray’s law and to achieve circular cross-section geometries. An alternative approach to addressing this tissue may be to use PU with a higher molecular weight which may achieve a semicrystalline structure that reduces rigidity, ensuring a smoother wire extraction^29^.

Furthermore, maintaining cross-sectional circularity within the hydrogel channels remains a significant challenge. We speculate that there are two possible causes of these irregularities: the spiral shape formed at bifurcations due to twisting can result in anisometric oval-shaped cross-sections after PU coating, and the twisting process may introduce gaps between wires at and near bifurcations. The use of smaller diameter template wires and tighter twisting potentially with mechanical assistance, may minimise these effects. Increasing the number of dips may also help to mitigate this issue, however as mentioned above, this may cause additional bead formation during dip coating and cracking at the hydrogel.

Finally, we fabricated seven-order bifurcating hydrogel channel networks using the methods developed here. While the modular nature of both rigs allows the orders to be easily increased, it does not match the full 20-30 orders of bifurcations necessary to model an entire human arterial vascular tree^17^. Further refinements to the method may be needed, for instance using automation to accelerate the wire template fabrication method if high fidelity models of the entire human vasculature are needed.

## 4. Conclusion

In this work we have put a ‘twist’ on our previous wire templating methods for generating multi-scale models of vasculature in vitro. This twisted wire templating method is an inexpensive process for generating perfusable higher order complex hierarchical networks that fractally branch and narrow at bifurcations within hydrogels. This approach is significantly faster and less labour intensive than previous techniques and enables the generation of higher order. This model has the potential to advance vascular research through being able to mimic organ level structure by creating systems branching from macro to micro scales.

## 5. Methodology

### 5.1 Wire spindle preparation

Copper wires (100 μm diameter, RS Components, UK) were cut to ∼32 cm long segments and wound onto spindles made up of electrical cable (diameter 1.5 mm, RS Components, UK) cut to 3 cm long segments to facilitate subsequent wire handling. The ends of each copper wire were fixed to spindles using 0.5 cm long segments of tape (RS Components, UK). The wire was then wound by rotating the spindle whilst maintaining tension on the wire. ∼0.5 cm of straight wire was intentionally left at the free end to facilitate later unwinding prior to twisting. This procedure was repeated until of the requisite number of wire spindles (up to 128) were prepared.

### 5.2 Wire bundling and organisation

To prevent wire tangling during the later processing such as dip-coating and wire twisting, wires wound onto spindles were grouped into bundles of four and systematically merged.

First, ∼10–12 cm of wire was unwound from each of four spindles. These four wires were grouped together by securing their distal ends onto a benchtop using tape. A second segment of tape was then applied near the spindle side to hold the four wires together as a bundle. The corresponding spindles were subsequently taped together after winding or unwinding the wires to equalise the wire lengths between the distal ends and the spools.

To further stabilise the distal fixation, two 4-wire bundles were combined into an 8-wire bundle by taping the wires together near the spindle side and securing the eight corresponding spindles together with tape. Pairs of 8-wire bundles were then secured at their distal ends onto the benchtop using tape, and the distal wire segments were twisted by approximately 4–5 turns. The same bundling procedure was repeated sequentially to form 16-, 32-, and 64-wire bundles, until bundles containing up to 128 wires were obtained.

This hierarchical organisation allowed the wire bundles corresponding to each bifurcation order to be easily accessed by sequentially removing the most recently applied tape.

### 5.3 Formation of seven-order bifurcations

Hierarchical bifurcations were fabricated by progressively dividing organised wire bundles into equal halves and twisting each resulting pair for four rotations at predefined nodes. This sequential approach is applicable to templates containing up to 128 wires. For a standard seven-order, 128-wire template, the initial bundle was secured to a workbench using tape, temporarily divided into two 64-wire bundles, twisted for four rotations, and recombined. The desired length of the straight, non-twisted base segment was measured, and the bundle was firmly secured at this precise location using a hardwood peg.

To generate the first bifurcation node, the bundle was again divided into two 64-wire bundles and twisted together for four rotations. Each 64-wire bundle was subsequently split into two 32-wire bundles, which were twisted for four rotations to form the distal branches before being recombined into 64-wire bundles. This symmetric division and twisting process was repeated at the next predefined segment length to create the second bifurcation. Throughout the procedure, tape constraints from preceding levels were progressively removed to expose the wires for the next split, while newly formed bundles were re-secured with tape to maintain spatial organisation.

To support higher-order bifurcations (orders three through seven), a temporary frame was constructed using interlocking plastic bricks and a baseplate. The height of frame was dynamically adjusted between 12 and 17 cm to accommodate the growing bifurcation order. The identical twist-bifurcation procedure was sequentially repeated at designated nodes until the final structure was complete. Post-fabrication, the wire spindles were secured to the support structure by sticking the spindle with the interlocking plastic bricks using tapes, so the frame was locked to the baseplate to prevent tangling during subsequent handling.

For templates originating with fewer than 128 wires, this exact fabrication strategy was preserved, with the maximum achievable branching order correspondingly reduced based on the initial bundle size.

### 5.4 Assembly and operation of dipper rig

To generate PU-coated bifurcating wire templates, a custom dipper rig was fabricated by laser cutting (Fusion Pro 36, Epilog Laser, UK) and subsequently assembled with the required electrical components. All computer-assisted design files for generating the dipper rig are provided in the electronic supporting information.

The dipper rig was first assembled and fixed where necessary using the laser-cut structural components. The installation and the connection of electrical components and the corresponding parts was previously described^12^.

For operation, the Arduino microcontroller was connected to a computer via USB after opening either the Arduino web editor or the desktop software. The control code provided in the electronic supporting information was copied into the sketch, verified, uploaded to the Arduino microcontroller, and then executed to control the dipping process^12^.

### 5.5 PU dip-coating of seven-order wire templates

The fully twisted wire template was transferred to the dipper rig for PU dip-coating. The parent bundle was threaded through the bottom hole of the rig, and the section before the first bifurcation was secured using tape and wrapped around the bottom beam of the frame. At the top of the rig, the wire spindles were inserted into the holes of the slide-in connectors and secured using tape to prevent loosening during coating.

After loading, the wire template was gently tensioned, and the spindles were rotated several turns to straighten the structure and minimise bead formation. The dipper rig was then assembled by connecting the frame to the connector supports and reinserting the linear rails into the bearings. 3.5L PU (Chase Corporation, USA) was poured into the container. A lid was placed on the rig to minimise airflow and prevent premature drying of the PU solution.

Dip-coating was performed using the optimised parameters identified in the preliminary experiments, with a dipping velocity of 1 mm s^−1^ and 15 dipping cycles. After coating, the template was allowed to air-dry for approximately 2 h at room temperature.

### 5.6 Silane coating of wire templates

After dip-coating, the wire template was detached from the dipper rig and subjected to silane coating. Prior to salinisation, the wire template was loaded to a hollow cylinder rig (the design file was provided in the electronic supporting information), and then plasma-treated using either oxygen plasma for 30 s ((PDC-001HP, Harrick Plasma, USA).

For silane coating preparation, Chloro(trimethyl)silane (1500 μL, Sigma-Aldrich, UK) was transferred into a glass vial using a glass syringe, which was then washed twice with Isopropyl alcohol (IPA, Sigma-Aldrich, UK). Before resealing the chloro(trimethyl)silane bottle, the headspace of the bottle was purged with nitrogen and the bottle was sealed with parafilm.

For vapour-phase salinisation, the inner wall of the vacuum chamber, including the sealing rubber, was wiped with tissue soaked in IPA. The the wire templates and the vial of chloro(trimethyl)silane were then placed inside the vacuum chamber. The chamber was sealed and connected to a nitrogen-driven vacuum line. Vacuum was applied until the pressure reached approximately −0.8 to −1.0 and stabilised, after which the chamber was closed, and the pump was switched off.

The chamber was left under vacuum for 30 min to allow evaporation of the chloro(trimethyl)silane. If liquid silane remained, the chamber was vented, the wire template position was adjusted if necessary, and vacuum was reapplied under the same conditions. This process was repeated in 30 min intervals until the chloro(trimethyl)silane had completely evaporated.

After coating, the wire templates were removed from the chamber. The vacuum chamber was wiped with IPA, and the glass container and vial were washed with IPA.

### 5.7 Alignment rig fabrication

To facilitate hydrogel casting around high-order wire templates and provide stable world-to-chip interfaces, custom-designed alignment rigs were fabricated. The capillary blocks with holes, capillary blocks without holes, pipette block, connector rails, and threading arms were manufactured by 3D printing, while the bottom board was fabricated by laser cutting. All computer-assisted design files and component lists for fabricating the alignment rig are provided in the electronic supporting information.

To reduce cracking at the outlet side during subsequent wire extraction, the connector rails (n = 4) were coated with polydopamine to improve bonding with polyacrylamide. The connector rails were first washed with acetone (>99%) and ethanol (>99%), then fully immersed in 2.0 mg mL^−1^ dopamine hydrochloride (Sigma-Aldrich, UK) prepared in 10 mM tris-hydrochloric acid buffer adjusted to pH 8.5 using 1 M hydrochloric acid (Sigma-Aldrich, UK) for 2 h at room temperature. Excess polydopamine was washed away with deionised water, and the coated parts were air-dried overnight at room temperature.

For preparation of the inlet pipette segment, a full 10 mL glass pipette was cut using a glass cutter to obtain a segment of approximately 3 cm in length. The cut edge was then fire-polished using a Bunsen burner (VWR, UK) to smooth the opening and remove sharp edges. For preparation of the outlet guide glass capillaries (2 mm outer diameter, World Precision Instruments, UK) was previously described^12^.

The outlet-side capillary block (8 × 16 × 10 mm) and inlet-side pipette block (16 × 16 × 10 mm) were then fabricated using the same PDMS-casting procedure. All holes on one side of each block were first sealed with tape. 16 guide glass capillaries were inserted into the outlet-side capillary block, while the pipette segment was inserted into the inlet-side pipette block. Sylgard 184 PDMS (Dow, UK) and curing agent were mixed at a 10:1 wt/wt ratio, and approximately 1 mL was poured into the middle reservoir of capillary and pipette blocks, respectively, to secure the inserted components. For capillary blocks without through-holes on the inlet side, PDMS was directly poured into the middle reservoir of the block. The blocks were cured at 65 °C for >2 h, after which the tape was removed. The embedded guide glass capillaries were retained for subsequent wire loading.

### 5.8 Wire loading to alignment rig

After dip-coating, the wire templates and their spindles were detached from the dipper rig. Additional wire length (10 cm) was unwound from each spindle to facilitate manipulation and straightening. Adjacent wires were first aligned and separated into bundles according to the intended bifurcation pattern. For the seven-order configuration, the wires were organised into 16 wire bundles in total. Each bundle was clamped at approximately 5 cm from the free end, twisted for approximately 4 cm, and trimmed to obtain even bundle lengths.

The twisted section of each wire bundle was then threaded into a guide glass capillary fixed within the capillary block. Bundle ends were secured onto the blocks by bending the end segments around the guide glass capillary edge. On the inlet side, the parent wire bundle was threaded into the pipette block.

After the wire bundles had been loaded through the capillary blocks, the bundles were carefully straightened from the glass capillary side of the rig and pushed inward by approximately 5 mm. Using magnification glasses and tweezers, the bundles were then separated sequentially according to the bifurcation order, beginning with the 4th-order bifurcation and continuing through the 5th to 7th orders, to restore the intended three-dimensional branching pattern prior to gel casting.

### 5.9 Assembly of alignment rig

The alignment rig was assembled by securing the male threading arm to the bottom board using an M3 × 8 mm screw (RS Components, UK) together with an M3 threaded insert (4 mm diameter, 4.78 mm depth, RS Components, UK). The connector rail was then attached to the threading arm. The wire-loaded capillary block and pipette block were connected to the alignment rig, taking care to maintain the organisation of the wire bundles and avoid tangling. The same procedure was repeated on the opposite side to attach the second connector rail and the female threading arm, which was also secured using an M3 × 8 mm screw and threaded insert.

After assembly, the pipette block and capillary block sides were separately sealed with tape to prevent disassembly during handling. Air-dry clay was then applied around the alignment rig to fill gaps between adjacent blocks, and the entire rig was wrapped with parafilm. Care was taken to ensure that no clay remained exposed. These sealing steps were necessary to minimise leakage of polyacrylamide from the assembled alignment rig during gel casting.

### 5.10 Microchannel formation by wire extraction

Microchannels were generated in polyacrylamide hydrogels using the loaded alignment rig. Before gel loading, deionised water was introduced into both the inlet pipette segment and outlet guide glass capillaries. A small air plug was then maintained in each capillary to separate the water phase from the subsequently loaded gel precursor and minimise mixing at the interface. The water–air interface was adjusted such that the final gel boundary was positioned approximately 3 mm from the end of each capillary or pipette segment.

After establishing the air plugs, 500 mL of polyacrylamide precursor solution was transferred into the reservoir of the alignment rig. Polyacrylamide precursor was prepared using acrylamide monomer (Sigma-Aldrich, UK) and N,N′-methylenebisacrylamide (bis-acrylamide, Sigma-Aldrich, UK) as the cross-linker. Acrylamide (5% w/v) and bis-acrylamide (0.3% w/v) were dissolved in deionised water. Polymerisation was initiated by addition of 1% (w/v) ammonium persulfate (APS, Sigma-Aldrich, UK) and accelerated by 0.1% (v/v) tetramethylethylenediamine (TEMED, Sigma-Aldrich, UK). After loading, the gel was allowed to polymerise for approximately 1 h at room temperature.

Gels were polymerised before gently extracting template wires at a rate of ∼1 cm/min by stringing the end of the wire using syringe pump. For bifurcating wire templates, extraction was performed from the parental side rather than the daughter side to minimise structural damage during removal. After wire extraction, the residual air gap within each capillary and pipette segment was removed.

### 5.11 FITC-dextran perfusion and imaging

A 2 mm guide glass capillary was connected to clear PVC tubing (1.5 mm inner diameter, 2 mm outer diameter; sourcing map; Amazon, UK), which was then attached to a 14G syringe needle. FITC-dextran solution (2000 kDa MW, 2.5 mg mL^−1^ in DI water; Sigma-Aldrich, UK) was loaded into a syringe and infused through the tubing-capillary setup until the solution reached the tip of the guide glass capillary, to remove any trapped air from the system. Once the fluidic pathway was free of air, the FITC-dextran solution was further infused into the inlet pipette segment.

After perfusion, the alignment rig was tilted toward the seventh-order bifurcation side to facilitate flow of the FITC-dextran solution into the smaller branches. This process was repeated until FITC-dextran solution was observed at the treated guide glass capillary outlet. Images were acquired using a Nikon Eclipse Ti2 microscope (Nikon, UK).

## Acknowledgements

This work was supported through a CRUK Multidisciplinary Award (DRCMDPA\100008), MRC Responsive Mode Grant (MR/Y000609/1), and MRC Gap Fund (UKRI236). We thank the CRUK Convergence Science Centre and the CRUK Microfabrication and Prototyping Facility for facility access. We thank Yuxin Zhang, Shusei Kawara for their assistance. We thank Miguel Hermida Ayala for training on confocal microscopy, Alisha Hunter for microfabrication training, and Paschal Egan & Tariq Malik for electronic device training. We are grateful to to Hamid Samivand and Hackspace Fellows for their assistance in part manufacturing.

## Conflict of Interest

The authors declare no conflict of interest.

## Author Contributions

J.D. and W.P. contributed equally to this work. J.D. contributed to the design of the 3D alignment rig and the final twisting method, and participated in manuscript preparation, dipping parameter optimisation, and the orthogonal test for microchannel fabrication. W.P. contributed to the design of the 3D dipper rig and participated in manuscript preparation, dipping parameter optimisation, and the orthogonal test for microchannel fabrication. F.A. contributed to the design of the alignment rig, manufacturing of both rigs, assisted with experiments, and participated in manuscript writing. H.T., L.B., and Y.X. contributed to the manufacturing of both rigs. B.C. provided logistical support and provided the conceptualization of the twisting method. S.H.A. was responsible for funding acquisition, idea generation, supervision, and manuscript writing. J.D., W.P., F.A., and S.H.A. contributed to edits.

## Notes

### Competing Interest Statement

The authors have declared no competing interest.

