## Supplementary material for "Micro-to-Macro Scale Hydrogel Microchannel Networks by Twisted Wire Templating": Supplemetary Information

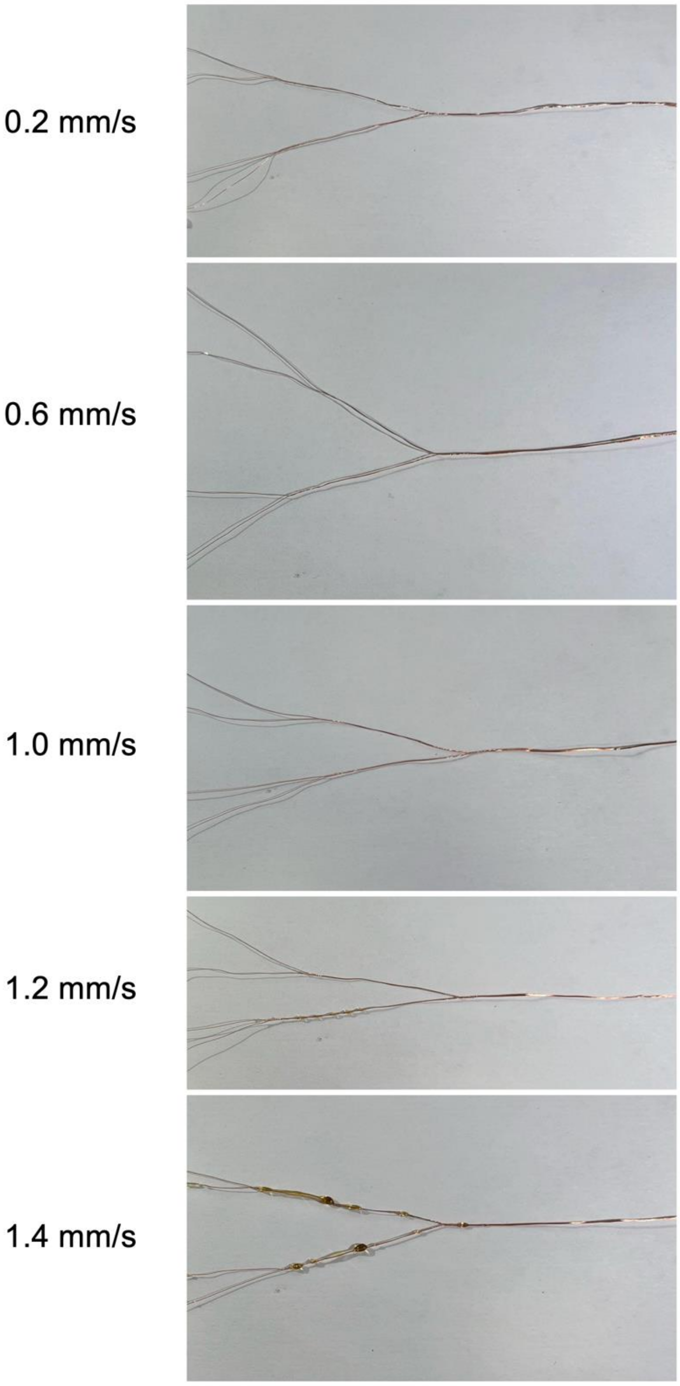


**Supplementary Figure 1. Representative results of the dipping velocity test using two-order bifurcating wire templates.** Wire templates were dip-coated for 10 cycles at velocities of 0.2, 0.6, 1.0, 1.2, and 1.4 mm $s^{-1}$. Coating morphology was compared across conditions to assess surface smoothness and bead formation. More pronounced bead formation was observed at higher dipping velocities, particularly at 1.4 mm $s^{-1}$.

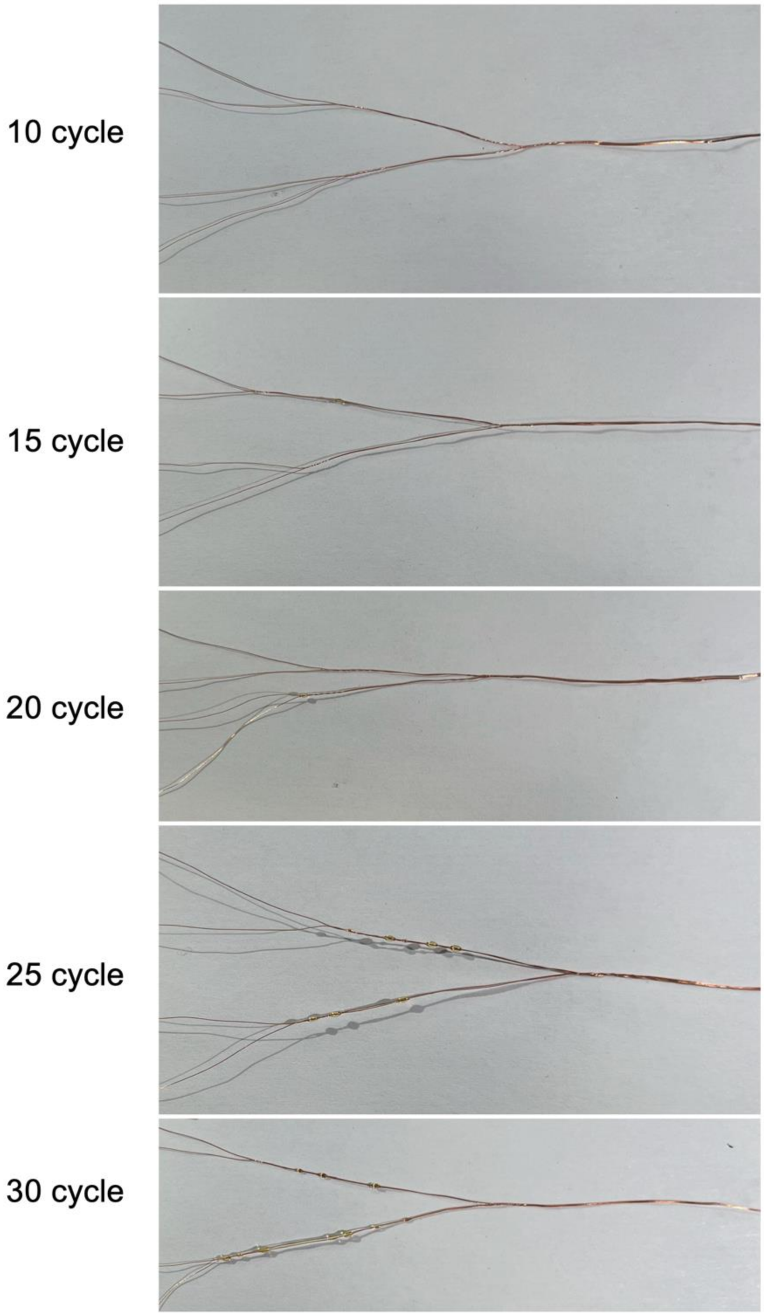


**Supplementary Figure 2. Representative results of the dipping cycle test using two-order bifurcating wire templates.** Wire templates were dip-coated at a constant velocity of 1 mm $s^{-1}$ using 10, 15, 20, 25, and 30 dipping cycles. Coating uniformity and bead formation were compared across conditions. More pronounced bead formation was observed at higher cycle numbers, particularly at 25 and 30 cycles.

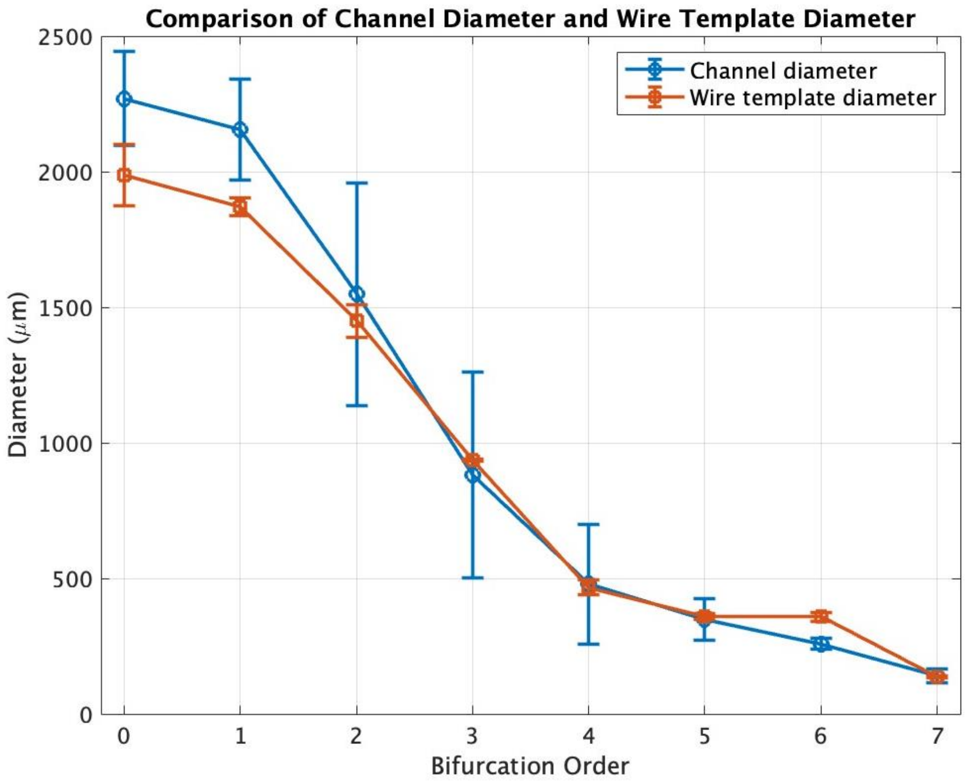


**Supplementary figure 3. Comparison of hydrogel channel diameter and wire template diameter across bifurcation orders.** Mean diameters of the wire templates and corresponding hydrogel channels are plotted as a function of bifurcation order (0th–7th order). Both structures exhibited a progressive reduction in diameter with increasing bifurcation order, indicating successful preservation of the hierarchical geometry during microchannel formation. Error bars represent standard deviation.

**Supplementary Table 1:** Optimisation factors for hydrogel channel formation at different levels

| Factors | | Level 1 | Level 2 |
|---|---|---|---|
| A | Capillary coating | No TMSPMA | With TMSPMA |
| B | Wire template coating | PU + TMSCl | PU only |
| C | Acrylamide concentration | 5% (w/v) | 7% (w/v) |

**Supplementary Table 2:** Fabrication setting combinations for seven-order multi-scale hydrogel channel according to Taguchi L4 ($2^3$) OA design

| Trials | Capillary coating (A) | Wire template coating (B) | Acrylamide concentration (C) |
|---|---|---|---|
| 1 | No TMSPMA | No silane | 5% (w/v) |
| 2 | No TMSPMA | With silane | 7% (w/v) |
| 3 | With TMSPMA | No silane | 7% (w/v) |
| 4 | With TMSPMA | With silane | 5% (w/v) |

**Supplementary Table 3**: Results of 3 response variables from 4 trials designed in Table S2

| Trials | Number of first types of cracks (#) | Rate of second types of cracks formation (#/sec) | Recovery (0) or breaking (1) condition of hydrogel (N/A) |
|---|---|---|---|
| 1 | 0 | 0 | 0 |
| 2 | 1 | 0 | 0 |
| 3 | 3 | 0.3625 | 1 |
| 4 | 2 | 0.2333 | 1 |
| Performance characteristics | Smaller-the-better | Smaller-the-better | Smaller-the-better |
